# Prophage induction mediated by quorum sensing signals alters soil bacterial community structure

**DOI:** 10.1101/805069

**Authors:** Xiaolong Liang, Regan E. Wagner, Bingxue Li, Ning Zhang, Mark Radosevich

## Abstract

Recent findings have revealed a role for bacterial quorum sensing communication in bacteriophage (phage) reproduction decisions. However quorum sensing based phage-host interactions remain largely unknown, with the mechanistic details revealed for only a few phage-host pairs and a dearth of information available at the microbial community level. Here we report on the specific action of individual quorum-sensing signals (acyl-homoserine lactones; AHLs varying in acyl-chain length from four to 14 carbon atoms) on prophage induction in soil microbial communities. AHL treatment significantly decreased the bacterial diversity (Shannon Index) but did not significantly impact species richness. Exposure to short chain-length AHLs resulted in a decrease in the abundance of different taxa than exposure to higher molecular weight AHLs. Each AHL targeted a different subset of bacterial taxa. Our observations demonstrate that individual AHLs trigger prophage induction in different bacterial taxa leading to changes in microbial community structure.

---

Bacteriophages (viruses of bacteria) may infect host cells via lytic and lysogenic cycles, both of which have shown ecological significance. For instance, lytic cycles of reproduction can impact population and community dynamics through lysis of host cells effectively re-routing dissolved organic carbon and other nutrients back to the dissolved pool, a process referred to as the “viral shunt”^1,2^. Lysogenic cycles in which the phage genome is inserted into the host cell genome without killing the host, may promote the host fitness and regulate metabolic functions through selective expression of certain phage encoded genes and transcriptional regulators without production of progeny phage particles^3,4^. Among the temperate phage, the mechanisms that control lysis-lysogeny decisions in natural environments remain unknown.

The “piggy-back-the-winner” (PtW) theory of phage-host population dynamics predicts that high microbial cell densities promote lytic to temperate (lysogenic) switching, highlighting the importance of lysogenic reproductive cycles at high host cell abundances^5^. Some microscopic counting-based examinations and viral metagenomic analyses provide evidence for PtW theory^5,6^. In contrast, the “kill-the-winner” (KtW) theory predicts that lytic infections are dominant and suppress the fastest growing hosts during times of high host cell densities, while lysogenic conversions are stimulated at low host cell abundances^7–9^. The long-standing KtW paradigm has also gained empirical support^10–12^. Both PtW and KtW suggest host-cell density may guide the viral reproductive strategies, although the paradigms propose contrasting fashions of host-cell density influences. Thus, cell density-dependent quorum sensing might have an important role in the lysogeny-lysis switch of temperate phages. Most recently, the molecular communication between viruses and between viruses and bacteria has shed light on the mechanism underpinning the phage lysogeny-lysis decisions in a few phage-host model systems^13–16^.

Quorum sensing for cell-density dependent communication among bacteria enables coordinated gene expression^17,18^. Quorum-sensing bacteria produce signaling molecules, such as different types of N-Acyl homoserine lactones (AHLs), with the concentration of released signaling molecules dependent upon bacterial population density^19–21^. Thus, quorum sensing plays a major role in adaptive survival and collective activity of bacterial communities. In an initial investigation evaluating the potential impact of QS on lysogeny-lysis switching, Ghosh et al.^22^ assessed the prophage induction response to exogenously added AHL mixtures of N- (butyl, heptanoyl, hexanoyl, ß-ketocaproyl, octanoyl, and tetradecanoyl) homoserine lactones and demonstrated that AHLs triggered viral production (i.e., switching from lysogenic to lytic viral reproduction) in soil and groundwater bacteria. The AHL-mediated prophage induction mechanism was demonstrated to be an SOS-independent process by using the single-gene knock-out mutation in the model system of *E. coli* with λ-prophage^22^. Similar studies by Silpe and Bassler^15,23^ revealed that the lysogeny-lysis switch of a *Vibrio* phage can be induced by the host-produced quorum-sensing autoinducers, in which the phage lysogeny-lysis decisions directly respond to host quorum-sensing molecular signals and cell density. However, the significance of this phenomena at the community level has not been investigated except for the previously noted report of Ghosh et al.^22^.

Communication among phages through phage-encoded arbitrium peptides was first described by Erez et al.^13^, and the following studies^14,24–26^ revealed the molecular basis for the production, detection, and consequences of the short signaling peptides on phages lysogeny-lysis decisions. Notably, these reports also showed that phages communicate only with their close relatives using a very specific arbitrium peptide, which suggests that phage communication peptides act in a taxon-specific manner just as bacterial quorum-sensing signals. Inspired by the above studies, especially the phage-bacterium quorum sensing connections, we hypothesized that any single quorum-sensing signal should only induce prophages within a small subset of closely related host bacteria. Towards that end, we tried to determine the impacts of the addition of individual AHL signaling molecules on prophage-induction and further assess the resulting impact of phage-mediated host cell lysis on bacterial community composition.

## Results

### Prophage induction

Soil samples from an agricultural field were collected near the end of winter (5 March) a time when potentially high proportions of lysogens exist in the bacterial community according to KtW. Bacteria were extracted, purified and concentrated from multiple soil samples in a way that eliminated most extracellular viruses. The resulting bacterial suspensions were pooled for use in prophage induction assays. The pooled cell concentrates had a bacterial count of 1.28 × 10^8^ cells per ml, and the density of viruses was below detection (approximately < 1 × 10^5^ particles ml^-1^). To determine the prophage-inducing effects of mitomycin C (MIT) and eight AHLs (refer to Table 1 and Supplementary Fig. S1 for molecular weight and structure) on bacterial populations, the induced samples were compared directly to the uninduced control samples based on viral/bacterial abundance and bacterial community structure.

**Table 1.** Information of the used inducing agents, quorum-sensing N-Acyl homoserine lactones (AHL1–8) and mitomycin C (MIT), in this study.

| Symbol | Name | Molecular weight | Formula |
| --- | --- | --- | --- |
| AHL1 | N-Butyryl-DL-homoserine lactone | 171 | C<sub>8</sub>H<sub>13</sub>NO<sub>3</sub> |
| AHL2 | N-Hexanoyl-DL-homoserine lactone | 199 | C<sub>10</sub>H<sub>17</sub>NO<sub>3</sub> |
| AHL3 | N-(β-Ketocaproyl)-L-homoserine lactone | 213 | C<sub>18</sub>H<sub>33</sub>NO<sub>3</sub> |
| AHL4 | N-Heptanoyl-L-homoserine lactone | 213 | C<sub>12</sub>H<sub>21</sub>NO<sub>3</sub> |
| AHL5 | N-Octanoyl-DL-homoserine lactone | 227 | C<sub>18</sub>H<sub>31</sub>NO<sub>4</sub> |
| AHL6 | N-(3-Oxododecanoyl)-L-homoserine lactone | 297 | C<sub>10</sub>H<sub>15</sub>NO<sub>4</sub> |
| AHL7 | N-Tetradecanoyl-DL-homoserine lactone | 311 | C<sub>16</sub>H<sub>27</sub>NO<sub>4</sub> |
| AHL8 | N-(3-Oxotetradecanoyl)-L-homoserine lactone | 325 | C<sub>11</sub>H<sub>19</sub>NO<sub>3</sub> |
| MIT | Mitomycin C | 334 | C<sub>15</sub>H<sub>18</sub>N<sub>4</sub>O<sub>5</sub> |

Viral and bacterial abundance was quantified to assess the induction of prophages in host cells. The MIT-treated samples had significantly lower bacterial abundance and notably higher viral abundance than the control samples (*P* < 0.01, t-test, Fig. 1). Though, no significant changes in bacterial abundance was observed in the eight AHL-treated samples compared to the control samples, relatively lower bacterial abundance was observed in AHL1-, AHL2-, AHL4-, and AHL6-treated samples. The viral abundance in all AHL-treated samples was higher than that in the control samples, and the difference of viral abundance in AHL1-, AHL2-, AHL5-, or AHL7-treated samples compared with the control samples was statistically significant (*P* < 0.05, t-test, Fig. 1b).

**Fig. 1.**
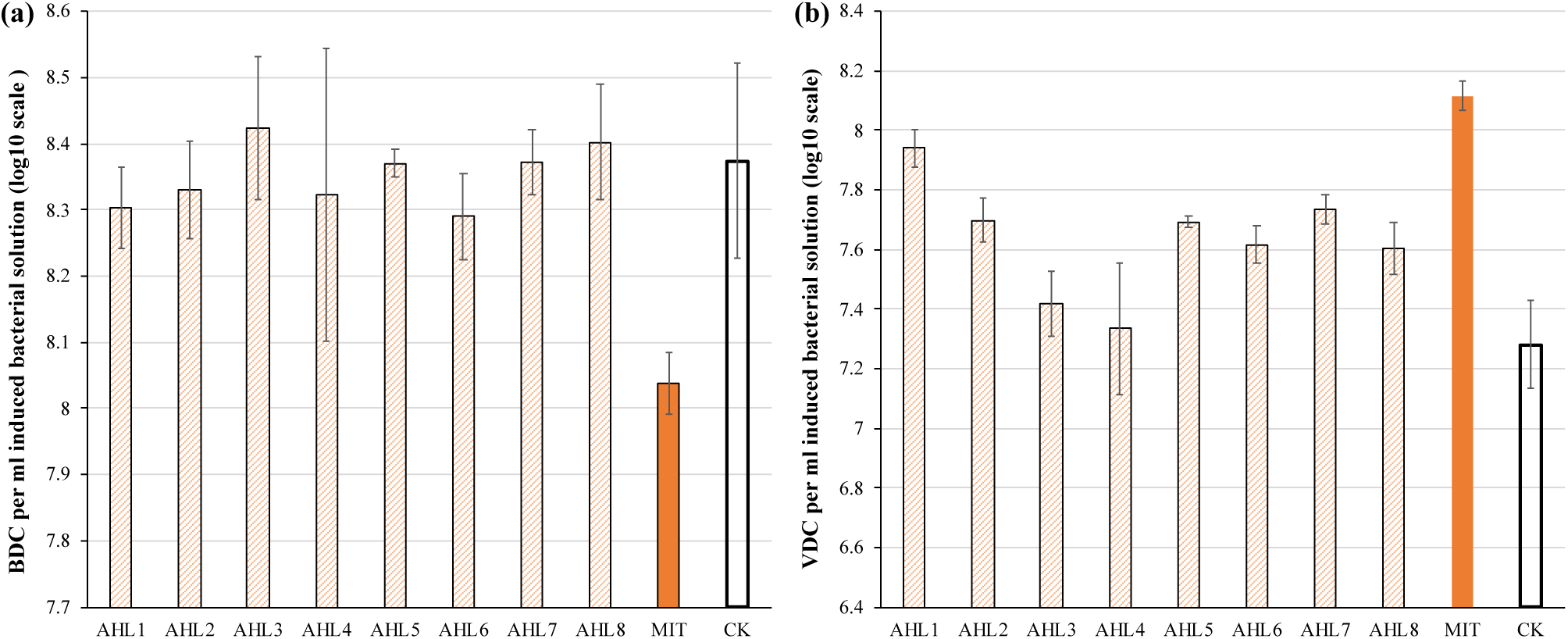
Direct counting of bacteria and viruses in the induced and control samples. Induction agents included mitomycin C (MIT) and N-(Butyryl, Hexanoyl, β-Ketocaproyl, Heptanoyl, Octanoyl, 3-Oxododecanoyl, Tetradecanoyl, and 3-Oxotetradecanoyl) homoserine lactones (represented by AHL1– 8). Each bar represents the mean value (n = 3), and the error bars show standard deviation.

### Bacterial community diversity and composition

To evaluate the impacts of prophage induction at community level, the bacterial species richness and community diversity were estimated based on species number and Shannon index, respectively as determined by analysis of 16S rRNA gene sequencing of un-lysed cells following the induction assays. The bacterial community in MIT-treated samples had significantly higher species richness and diversity than that in the control samples (*P* < 0.01, t-test, Fig. 2). Contrastingly, the bacterial species diversity in AHLs-treated samples was lower than that in the control samples (*P* < 0.05, t-test, Fig. 2a). Similar results were obtained for inverse Simpson index (data not shown). There was no statistical difference in bacterial species richness between each AHL treatment and the control samples (Fig. 2b).

**Fig. 2.**
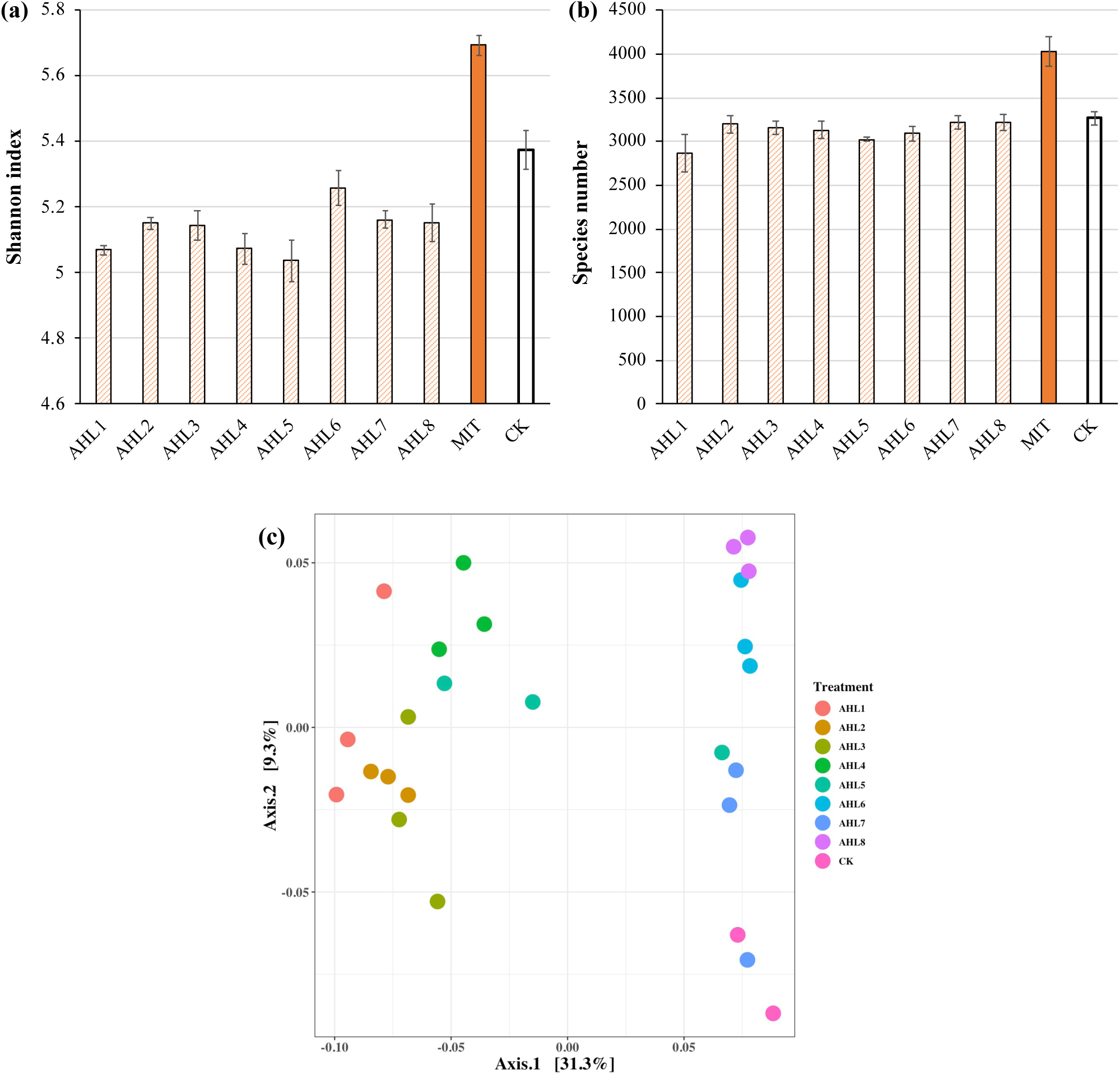
Alpha- and beta-diversity of bacterial communities. a) Shannon index of bacterial communities. b) Bacterial species richness. Data was shown as the mean value of triplicate samples, and the error bars show standard deviation. c) Principal coordinate analysis (PCoA) of bacterial community composition. Samples were color-coded based on treatment of AHLs and the control (CK). AHL1–8 stands for induction assay of N-(Butyryl, Hexanoyl, β-Ketocaproyl, Heptanoyl, Octanoyl, 3-Oxododecanoyl, Tetradecanoyl, and 3-Oxotetradecanoyl) homoserine lactones, respectively.

The principal coordinate analysis (PCoA) was used to visualize the dissimilarities of bacterial community structure between samples after prophage induction and showed that the communities were separated according to the specific AHL signal added to the bacterial extracts (Fig. 2c). The bacterial community structure in the induction assays using AHL1–5 were notably different from those in the control samples. Some dissimilarity was also observed in bacterial community structure among samples treated with different AHLs. For example, communities resulting from treatment with AHL1-5 were distinctly different from communities treated with AHL6-8 (Fig. 2c). MIT-treated samples were clearly separated from controls and AHL-treated samples, and the first two principal coordinates explained 60.4% of the variation of bacterial community structure (Supplementary Fig. S2).

### Bacterial taxonomic profiles following prophage induction

To further examine the differences in bacterial community structure resulting from AHL-mediated prophage induction, we compared the abundance of each bacterial taxonomic group in every induction treatment to that in the control samples at both class and genus level. AHL1 treatment led to a decreased abundance of the greatest number of taxa including: Actinobacteria, Alpha-Proteobacteria and Acidobacteria Gp6 at the class level (Fig. 3) and *Microbacterium, Actinophytocola, Mamoricola, Nocardioides, Novosphingobium, Lysobacter, Pandoraea* and *Sphingomonas* at the genus level (Fig. 4). AHL2 treatment led to a decreased abundance of two bacterial classes, Actinobacteria and Alpha-Proteobacteria, and genera, *Brevundimonas, Mesorhizobium, Lysobacter* and *Sphingomonas* (Fig. 4). AHL5 treatment caused a decreased abundance of the second greatest number of bacterial taxa including Acidobacteria Gp6, Gp17 and Actinobacteria at the class level and *Mamoricola, Gp17, Bosea, Nocardioides, Bradyrhizobium, Novosphingobium* and *Gp6* at the genus level (Fig. 4). Treatment with AHL3 and 4 resulted in a decreased abundance of only one class, Chlamydiae. However, AHL3 and 4 had distinct induction effects at the genus level where the abundance of *Streptomyces, and unclassified Anaerolinaceae and Parachlamydiaceae* declined with AHL3 treatment and *Bosea, Verrucomicrbium* and *Pedobacter* in AHL4 exposure. The AHLs discussed above also resulted in an increased abundance of some bacterial taxa, *e.g.*, classes of Gamma-Proteobacteria, Bacilli and Flavobacteria although the positive affect on some taxa may be indirectly attributed to prophage induction. It is interesting to note that an increased abundance of *Pseudomonas* was observed in the treatment with AHL1–5 that had significant shifts in the density of specific bacterial groups. The bacterial genera having increased abundance after treatment with some AHLs also included *Filimonas, Cellvibiro, Duganella, Pelomonas* and *Flavobacterium* and certain other unclassified genera. The bacterial taxonomic profiles in the treatment of AHL6, 7 and 8 had no statistical difference compared with the control samples.

**Fig. 3.**
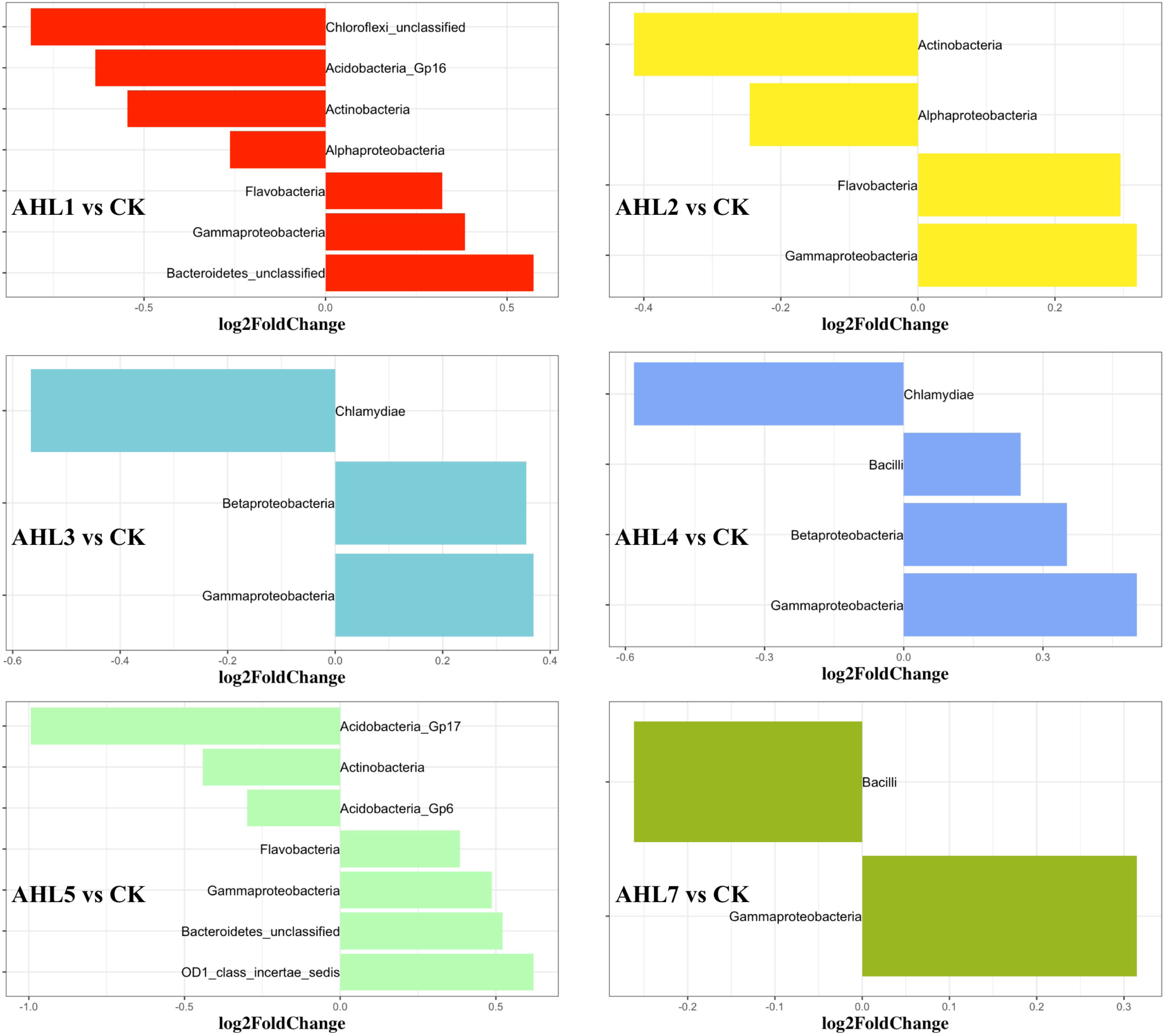
Class-level differences of the bacterial community composition between each induction assay of N-(Butyryl, Hexanoyl, β-Ketocaproyl, Heptanoyl, Octanoyl, 3-Oxododecanoyl, Tetradecanoyl, and 3-Oxotetradecanoyl) homoserine lactones (shown as AHL1–8, respectively) and the control samples. Only statistically significant differences (*P* < 0.01) were shown. The direction of bars represents lower (left) or higher (right) abundance of the specific bacterial taxonomic group in the induction assay.

**Fig. 4.**
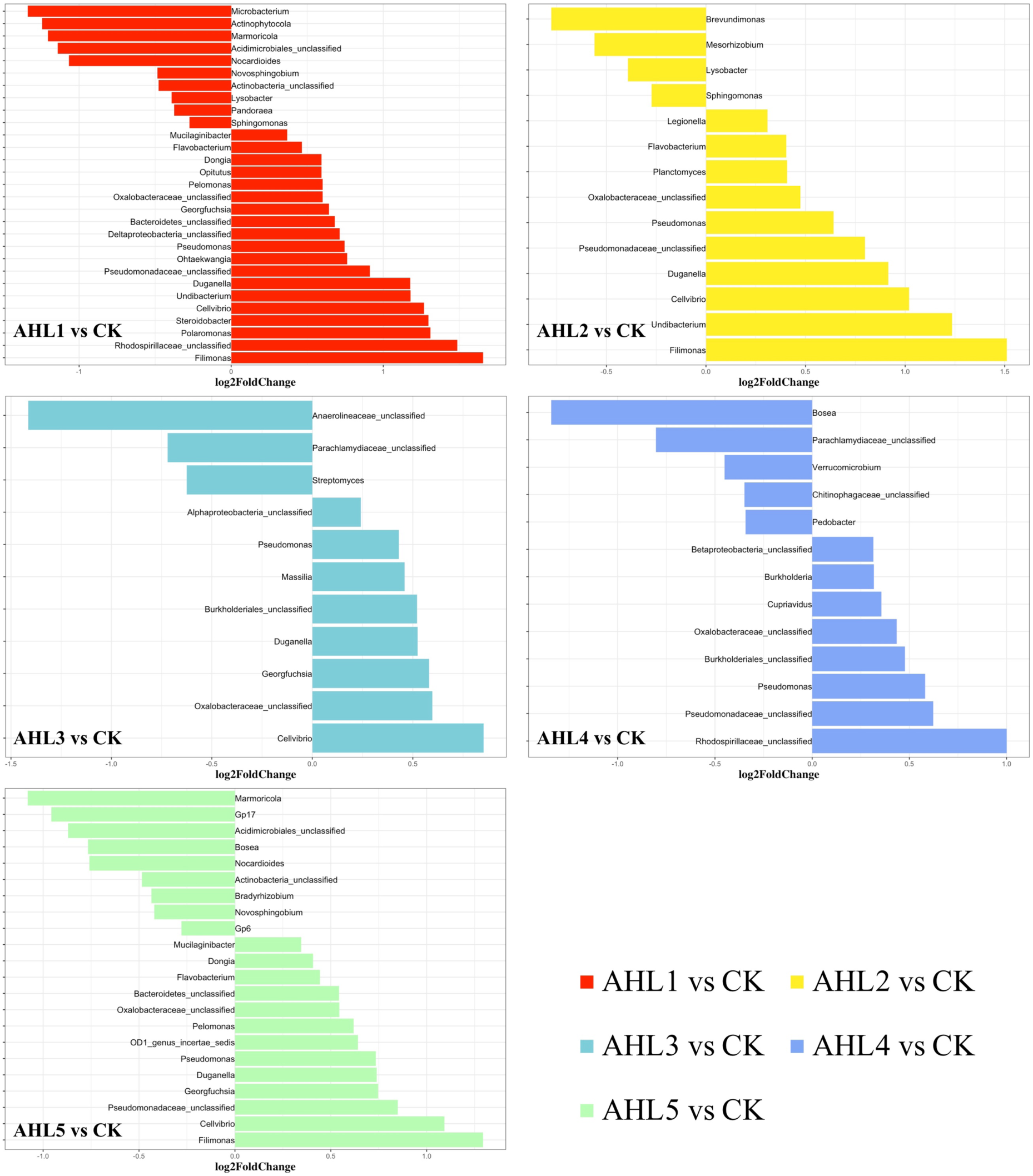
Genus-level differences of the bacterial community composition between each induction assay of N-(Butyryl, Hexanoyl, β-Ketocaproyl, Heptanoyl, Octanoyl, 3-Oxododecanoyl, Tetradecanoyl, and 3-Oxotetradecanoyl) homoserine lactones (indicated by AHL1–8, respectively) and the control samples. Only statistically significant differences (*P* < 0.01) were shown. The direction of bars represents lower (left) or higher (right) abundance of the specific bacterial taxonomic group in the induction assay.

MIT treatment stimulated much broader impacts on the taxonomic profiles of the bacterial community than the AHLs suggesting that the prophage induction response brought about by exposure to MIT was less specific than any of the AHLs used as inducing agents. The abundance of six classified bacterial classes, including Flavobacteria, Bacilli, Gamma-Proteobacteria, Acidobacteria Gp7, Opitutae and Verrucomicrobiae, declined in MIT treatment (Fig. 5a). Up to 53 genus-level bacterial groups, such as *Aeromonas, Flavobacterium, Albidiferax, Chitinimonas, Arthrobacter, Chryseobacterium* and *Paenibacillus*, had decreased density after MIT treatment (Fig. 5b). *Cellvibiro, Filimonas, Pelomonas, Flavobacterium* and *Pseudomonas*, the bacterial genera that had increased density after the treatment of some AHLs, had decreased density in the MIT treatment. The common bacterial genera having decreased density and shared by MIT and some AHL treatments included *Novosphingobium* and *Pedobacter*. MIT exposure also drew a lightly increased proportion of 44 bacterial genera (Fig. 5b).

**Fig. 5.**
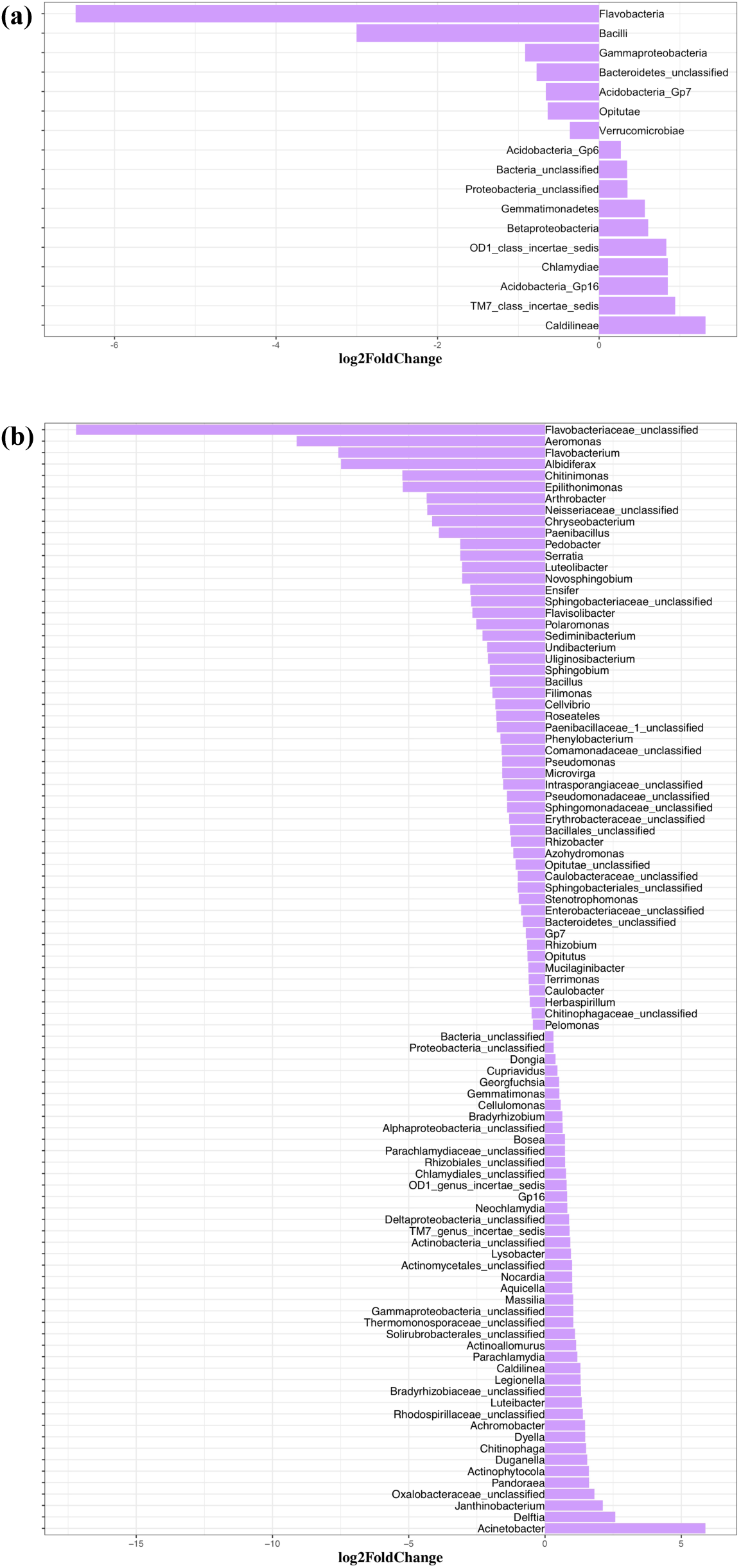
Systematic differences of the bacterial taxonomic composition between induction assay of mitomycin C and the control samples at Class (a) and Genus(b) levels. Only statistically significant differences (*P* < 0.01) were shown. The direction of bars represents lower (left) or higher (right) abundance of the specific bacterial taxonomic group in the induction assay.

## Discussion

This study revealed the lysogeny-lysis switch of some temperate phages is responsive to quorum-sensing autoinducers. The molecular basis of a host quorum-sensing autoinducer controlling a phage lysogeny-lysis decision has been recently characterized^15^. Subsequent studies reported phage responses to other types of host autoinducers^23,27^. Bacteria communicate only with their close relatives using specific quorum-sensing signals, thus phage-bacterium quorum sensing connections may also be molecular structure dependent. Building on this idea, we hypothesized that any single quorum-sensing signal should only induce prophages within a small subset of closely related host bacteria. In this study, eight AHLs of varying molecular weight and structure were selected for evaluation of phage responses of lysogeny-lysis switching and its significance in structuring the exposed bacterial communities.

Significant differences in viral abundance were observed in bacterial communities exposed to AHL1, 2, 5 and 7 compared with the control samples, consistent with a burst of viral production from prophage induction, especially in relatively specific host taxa, consistent with our hypothesis. A significant decrease in total cell abundance was typically not observed upon exposure to AHLs. This too is consistent with the hypothesis if each AHL triggered prophage induction in a relatively narrow range of perhaps less numerically abundant taxa.

However, even a small collection of lysogenic bacterial taxa triggered to enter the lytic cycle could result in changes in taxonomic profiles. We inspected the bacterial community diversity based on species richness and Shannon index. Significantly lower Shannon indices but no significant changes in species richness were observed after AHL treatment. These results suggest that AHL treatment decreased species evenness potentially by suppressing a subset of bacterial species and allowing dominance of some other species. In contrast, MIT treatment, as a broad-spectrum inducing agent, also possessing broad toxicity, commonly used for prophage induction, prompted an increase in both species richness and diversity relative to untreated controls. MIT treatment brought about a significant increase in viral abundance (*P* = 0.001) and notable decrease in bacterial abundance (*P* = 0.007) through a wide range of viral lysis, thus contributing to higher richness and evenness in bacterial taxonomic profiles.

To further examine the putative effects of AHL-dependent prophage induction on bacterial community structure, we determined changes in abundance (expressed as log2fold changes relative to control cultures) of the affected taxa. Up to 10 bacterial genera decreased in abundance after any single AHL treatment, and a total of 23 different genera for all AHL treatments combined. Decreased abundance of *Lysobacter, Novosphingobium, Sphingomonas, Bosea*, and *Nocardioides* was observed in at least two AHL treatments. The significant decrease in density of the AHL-targeted bacterial genera, e.g., *Lysobacter, Novosphingobium*, and *Sphingomonas*, suggests AHL-directed transition of prophages from lysogeny to lysis in these bacteria.

Quorum sensing has been demonstrated to be widespread among bacteria^28–30^, however AHLs have not been shown as quorum sensing signals for many of the AHL-impacted bacterial groups in the present study, such as *Nocardioides, Streptomyces, Pedobacter*, and *Verrucomicrbium*. Thus, we propose a new hypothesis that prophages in a host cell can respond to the bacterial quorum sensing signals of different taxa other than the infected host.

MIT treatment resulted in decreased abundance of 53 bacterial genera. Though the observed decreases may have resulted from virus-mediated host cell lysis upon prophage induction or simply from direct toxicity of the MIT, these broad-spectrum changes clearly contributed to the observed increases in evenness of bacterial species profiles and thus increases in the community diversity. Increased proportion of specific bacterial groups observed in AHL or MIT treatment might be derived from competitive release^31^. Growth of some rare bacterial species might also be promoted which contributed to the increased species richness in MIT treatment. Susceptible bacterial species were lysed by chemical induction of prophages allowing the remaining competitors to utilize the resources more fully, and the remaining members of the bacterial community may also have access to the nutrients released through viral shunt^1,32^.

Lysogeny has been demonstrated to be widespread and a common viral life strategy in nature and shown to have links with dynamics of nutrient regime and host density^33^. Chemical induction assays, like MIT, have been adopted to assess lysogeny among viral and bacterial communities^22,34,35^. Though the phage-bacterium connections through quorum sensing was recently discovered, the quorum sensing based prophage-induction are largely unknown except for a few phage-host pairs none of which were derived from soil, with little known about the influence of prophage induction on microbial community dynamics. In this study, we revealed that a vast range of bacterial taxonomic groups are putatively susceptible to prophage induction by host autoinducers or other chemical agents, and we also demonstrated how transitions from lysogeny to lysis of temperate phages responsive to different host autoinducers can have pivotal roles in influencing the bacterial taxonomic profile development and shaping the bacterial community structure. This research provides theoretical and methodological foundation for future study of phage-bacterium communication and the lysogeny-lysis switch of soil viral communities. For future study, the results reported here should be confirmed by including analysis of lysed bacteria as template for 16S rRNA sequence analyses. In addition, metagenomic and metatranscriptomic analyses before during and after induction assays may reveal more specific and direct evidence supporting QS-controlled lysogeny-lysis switching and the hypothesis addressed in this study may resolve more statistically robust relationships and provide unique high-resolution views of virosphere responses to host autoinducers.

## Methods

### Sample collection and bacterial extraction

Soil samples were collected from an agricultural field at the East Tennessee Agricultural Research and Education Center (Latitude = 35.899166; Longitude = –83.961120) on March 5, 2019. The extraction of bacterial cells from soils was performed as described elsewhere^36^ except a larger quantity of soil (300 g) was extracted to secure sufficient microbial biomass to complete all the induction assays described below. The soil to extraction buffer ratio was maintained as previously described^36^. The extracted bacteria were concentrated by centrifuging the slurries on a cushion of Nycodenz at 4000 g for 20 min at 4 °C. Supernatant including the nycodenz phase that contains the bacterial cells was transferred to 50 ml centrifuge tubes and was then centrifuged at 5,000 g for 20 min. The bacterial pellets were washed twice using sterile extraction buffer and resuspended in autoclaved 0.2 µm-filtered soil extract^22^ after decanting the supernatant to discard extracellular viruses. The bacterial extracts were pooled and homogenized by gently shaking prior being added to induction assay tubes containing the respective inducing agents or controls. Sterile soil extract was used as the medium for induction assays to provide native growth substrates and trace elements to support bacterial metabolism and viral reproduction in induced lysogenic cells. The bacteria and viruses were enumerated after incubation via epifluorescence microscopy direct counting as described^36^.

### In vitro prophage induction with AHLs as inducing agents

Eight different AHLs, (N-butyryl-, hexanoyl-, β-ketocaproyl-, heptanoyl-, octanoyl-, 3-oxododecanoyl-, tetradecanoyl-, and 3-oxotetradecanoyl-homoserine lactones)^21^ varying in molecular weight and structure (Supplementary Fig. S1) and MIT were selected for induction assays. The selected AHLs were designated AHL1 to 8 based on their molecular weight from lowest to highest (acyl C chain length refer to Table 1). Each AHL was dissolved in ethyl acetate acidified with acetic acid (0.1%, vol/vol) as stock solution and applied at a final concentration of 1 μM in the bacterial suspension. The required amount of each AHL compound in stock solution was added into glass tubes, and the tubes were gently shaken for evaporation of the solvents so that the AHL compounds bonded to the tube bottom as a film^22^.

The pooled bacterial extract was distributed into AHLs- and mitomycin C-coated glass tubes, each containing a 10 ml aliquot of bacterial suspension. Ten ml of the exact same of bacterial suspension was also distributed to clean glass tubes lacking any inducing agent to serve as control. Each treatment and control were prepared in triplicate. All suspensions were incubated in the dark for 18 h at room temperature^22^. The viruses and bacteria in the suspensions were enumerated using epifluorescence microscopy to determine the induction response due to each inducing agent. Viral and bacterial abundance in each sample was estimated by epifluorescence microscopy enumeration as previously described^37,38^.

### Bacterial 16S rRNA genes sequencing and statistical analysis

After 18 h incubation, 1 ml of each cell suspension from the induction assays was transferred to a new sterile centrifuge tube and treated with DNase I for 20 min. The reaction was terminated by addition of EDTA prior to centrifugation at 5000 g for 20 min at 4 °C. The bacterial pellets were washed twice with sterilized extraction buffer to remove all lysed bacterial cells and any residual undigested free DNA. The genomic DNA of un-lysed bacterial cells that survived prophage induction from each sample was extracted using PowerLyser PowerSoil DNA isolation kit (Qiagen, Hilden, Germany) and quantified with Nanodrop one^C^ spectrophotometer (Thermo Scientific). The DNA samples were sent to Genomics Core Laboratory at University of Tennessee (Knoxville, TN, USA) for sequencing. The V3-V4 region of 16S rRNA genes were amplified using PCR primer set (341F_CCTACGGGNGGCWGCAG, and 785R_GACTACHVGGGTATCTAATCC) for library construction, and finally sequenced via 300PE (paired-end) on the Illumina MiSeq platform (Illumina, USA) by using the manufacturers’ protocol.

The raw 16S rRNA gene sequence data with a total of 3,821,672 sequence reads was processed using the MOTHUR v.1.40.0 pipeline according to the MiSeq SOP^39^. Statistical analyses of the processed sequence data were performed using software R version 3.6.1 packages phyloseq^40^, vegan (version 2.5-2^41^), ggplot2^42^, and DESeq2^43^. Bacterial taxonomic composition and alpha-diversity were calculated, and beta-diversity was also assessed based on Bray-Curtis dissimilarity matrix. The quantification and statistical inference of systematic differences of the bacterial taxonomic composition between each induction assay and the control samples were performed using the package DESeq2.

## Data availability

The files of raw sequences were archived at the National Center for Biotechnology Information Databases (Sequence Read Archive) and can be obtained under project number PRJNA575893.

## Acknowledgements

This work was supported by a United States Department of Agriculture grant to MR (award number: 2018-67019-27792). Financial support from China Scholarship Council was also awarded to XL. The authors also acknowledge the constructive comments provided by anonymous reviewers that were helpful in improving the manuscript.

## Author contributions

X.L. and M.R. conceived and designed the study. X.L., R.E.W., B.L. and N.Z. collected and processed the field samples and conducted the experiments. X.L. completed sequencing and bioinformatics and worked with M.R. on data analyses. X.L. composed the manuscript, and all authors contributed to the revisions of the manuscript.

