## Supplementary information for "Prophage induction mediated by quorum sensing signals alters soil bacterial community structure"

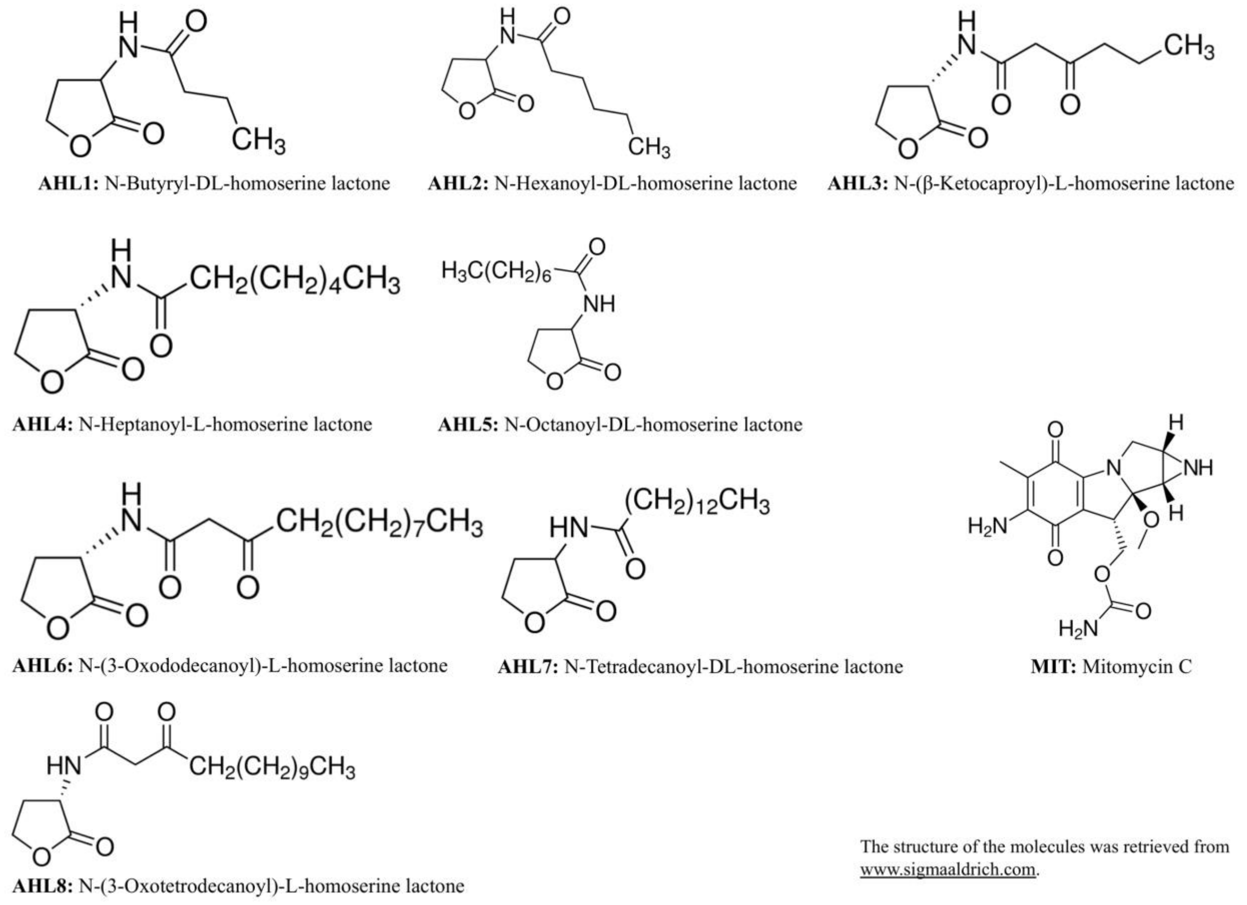


**Figure S1.** The molecular structure of quorum-sensing N-Acyl homoserine lactones (AHL1–8) and mitomycin C (MIT).

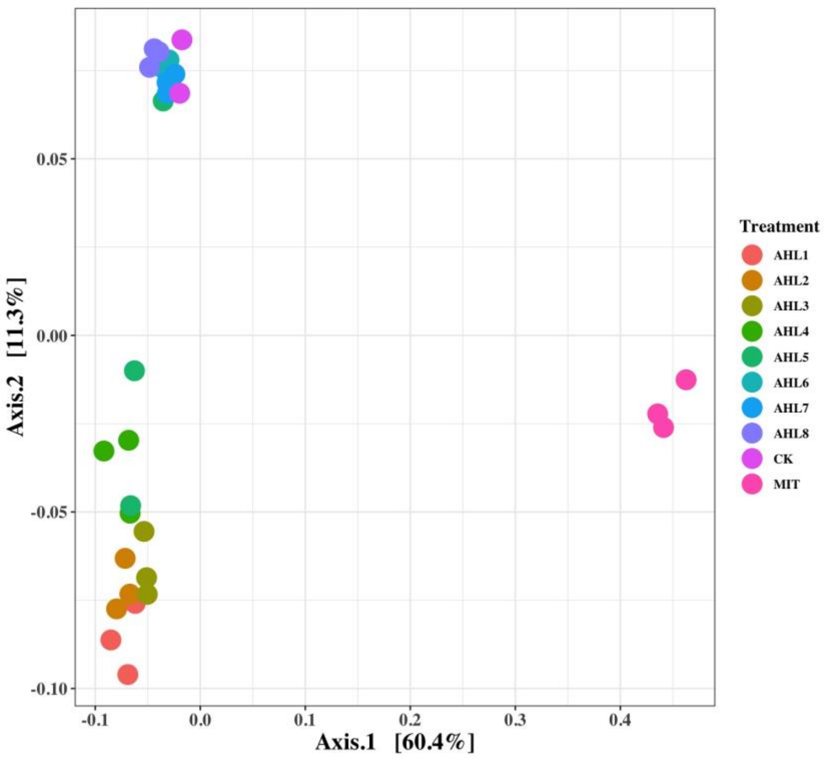


**Fig. S2** Principal coordinate analysis (PCoA) of bacterial community composition. Treatments, including induction assays of mitomycin C (MIT), N-(Butyryl, Hexanoyl, β-Ketocaproyl, Heptanoyl, Octanoyl, 3-Oxododecanoyl, Tetradecanoyl, and 3-Oxotetradecanoyl) homoserine lactones (represented by AHL1–8, respectively), and the control (CK) are color-coded.

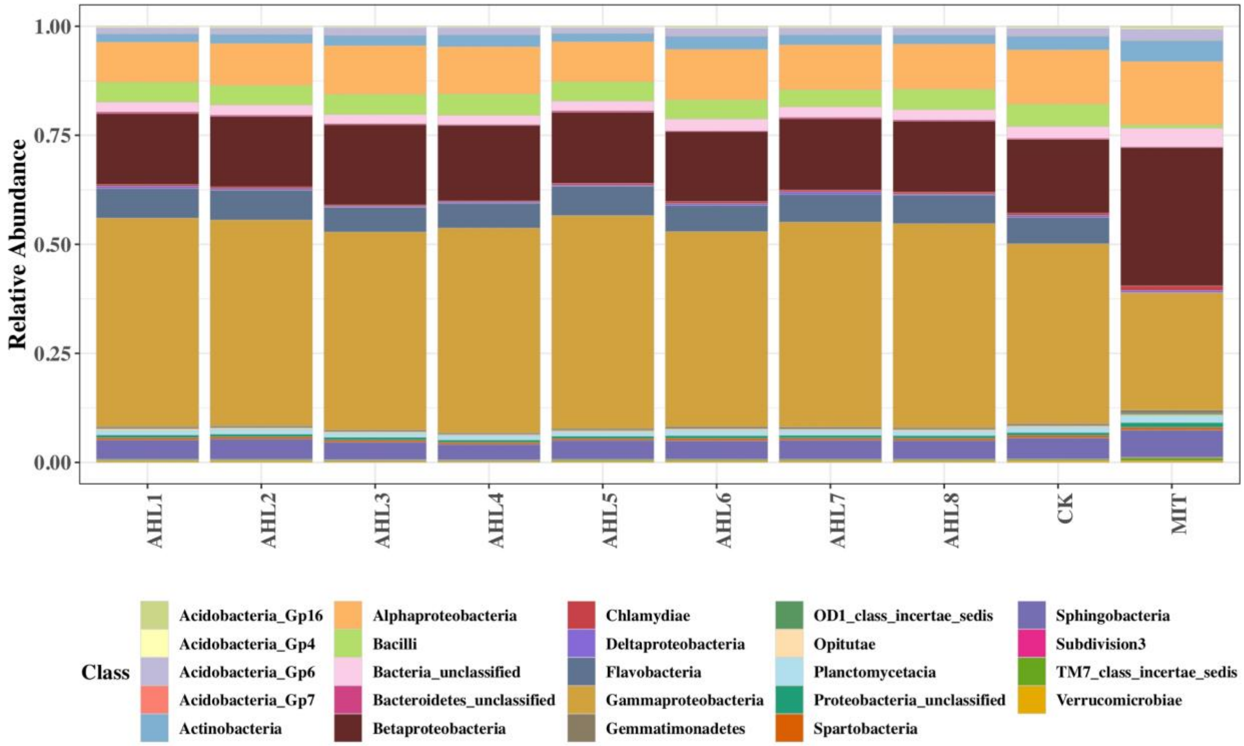


**Fig. S3** Bacterial community composition at Class level. Treatments include induction assays of mitomycin C (MIT), N-(Butyryl, Hexanoyl, β-Ketocaproyl, Heptanoyl, Octanoyl, 3-Oxododecanoyl, Tetradecanoyl, and 3-Oxotetradecanoyl) homoserine lactones (represented with AHL1–8, respectively), and the control (CK).
